# Colorectal adenoma presence is associated with decreased menaquinone pathway functions in the gut microbiome of patients undergoing routine colonoscopy

**DOI:** 10.1101/2025.07.29.667347

**Authors:** Ilona Vilkoite, Ivars Silamiķelis, Jānis Kloviņš, Ivars Tolmanis, Aivars Lejnieks, Elīna Runce, Krista Cēbere, Ksenija Margole, Olga Sjomina, Laila Silamiķele

**Author notes:** Ilona Vilkoite is a corresponding author [, Upeņu str, Rīga, LV-1084].

## Abstract

**Background:** Colorectal cancer (CRC) is among the most common malignancies worldwide, with colorectal adenomas recognized as well-established precursors to CRC. Changes in gut microbiota appear to be linked to CRC by promoting chronic inflammation, immune dysfunction, and metabolic issues that drive tumor growth and progression.

**Objectives:** To explore the relationship between gut microbiome composition and the presence of colorectal adenomas in patients undergoing routine colonoscopy.

**Materials and methods:** Patients were selected from those receiving standard colonoscopy based on strict inclusion and exclusion criteria, to minimize potential confounding factors such as previous colorectal surgeries, inflammatory bowel diseases, and the use of antibiotics or probiotics. Fecal samples were collected before bowel preparation for the colonoscopy procedure, and metagenomic shotgun sequencing was used to analyze the composition and functions of the gut microbiome.

**Results:** Overall, 136 participants were recruited, and 56 of them had colorectal adenomas. Although no distinction was observed in alpha diversity, beta diversity analysis indicated significant differences between the adenoma-positive and adenoma-negative groups. Signs of dysbiosis were found in patients with adenomas: increased abundance of the genera *Bacteroides* and *Prevotella* and decreased abundance of *Faecalibacterium* and *Anaerostipes* species. Beta diversity analysis showed statistically significant differences in the structure of the microbiota. Significant differences in the relative abundance of *UBA7597 sp003448195* were observed between groups. Functionally, decreased vitamin K2, SCFA (propionate) synthesis, along with lower Stickland fermentation activity was observed, indicating altered microbial metabolism. These changes may compromise epithelial barrier support, anti-inflammatory signaling, and energy metabolism in the colon.

**Conclusions:** The discovery of microbial taxa and functional pathways associated with the presence of adenomas underscores the potential of microbiota-based biomarkers and therapeutic strategies in the prevention and management of colorectal cancer.

## Introduction

Colorectal cancer (CRC) remains a significant global health challenge, representing a substantial burden on both healthcare systems and affected individuals (1). Owing to its high incidence and mortality rates, CRC ranks among the most prevalent and deadly malignancies worldwide. According to recent epidemiological data, CRC accounts for approximately 10% of all cancer cases and is the third most commonly diagnosed cancer globally, with an estimated 1.8 million new cases diagnosed annually (1).

The etiology of CRC is multifaceted, involving a complex interplay of genetic, environmental, and lifestyle factors (2). Major risk factors include high consumption of red meat coupled with low fiber intake, obesity, lack of physical activity, substance abuse, and persistent bowel inflammation (3). Among these factors, emerging research has increasingly recognized the pivotal role of the gut microbiome in the development and progression of CRC (4–6). The human colon harbors a diverse and dynamic community of microorganisms, collectively known as the gut microbiota, which plays a fundamental role in maintaining intestinal homeostasis and influencing various aspects of host physiology (7). Typically, polyps progress to malignancy following a well-defined path known as the adenoma-carcinoma sequence (8). Alternatively, 15 to 30% of CRCs develop through the serrated pathway (9).

Polyps in the premalignant stage from both pathways can be screened and removed during colonoscopy to prevent the formation of CRC. However, if polyps are incompletely removed or go undetected, this can lead to the emergence of interval cancers. The formation of colorectal polyps precedes the development of cancer and is impacted by a range of environmental factors and the host’s genetics (8).

Alterations in the composition and function of the colon microbiota, including, but not limited to, dysbiosis have been implicated in the pathogenesis of CRC, particularly when these changes disrupt immune homeostasis, barrier integrity, or metabolic balance. Mounting evidence suggests that dysbiotic changes in the gut microbiota can contribute to chronic inflammation, aberrant immune responses, and metabolic dysregulation within the colonic microenvironment, thereby promoting tumorigenesis and tumor progression (10). Understanding the intricate relationship between the colon microbiota and CRC pathogenesis holds immense promise for developing novel preventive and therapeutic strategies for this devastating disease.

Growing evidence suggests that qualitative or quantitative changes in the abundance of specific gut microbiota members could serve as markers for the future development of colorectal neoplasia. Although various studies have explored changes in gut microbiota composition in the context of colorectal adenomas, the results remain inconclusive (11–13), highlighting the need for further research. Investigations across diverse populations offer additional insights.

Our study aimed to evaluate the complex interplay between changes in the composition of the gut microbiota and the development of colorectal adenomas, known as precursors of CRC. The primary hypothesis was that the gut microbiota composition differs between individuals with and without colorectal adenomas. We described observed differences in gut microbiota composition and functions between individuals with and without colorectal adenomas.

## Materials and methods

This was a single-centre case-control study conducted by a single expert with an adenoma detection rate (ADR) of 36% in the screening population. The study was conducted from April 1, 2021, to April 22, 2022, at the outpatient endoscopy unit of Health Center 4 in Riga, Latvia, as well as the Academic Histology Laboratory and the Latvian Biomedical Research and Study Center.

### Informed consent and ethics statement

Written informed consent was obtained from all participants. All procedures were performed in accordance with the ethical standards of the Declaration of Helsinki. The study protocol and patients’ recruitment were approved by the Central Medical Ethics Committee of Latvia (Permit No. 01-29.1.2/1751).

### Participants and study design

In total, 146 patients were recruited for the study; however, ten patients were excluded due to poor bowel preparation, ulcerative colitis, or colorectal cancer. Overall, 136 patients over the age of 18, undergoing colonoscopy for various reasons, who provided informed consent and met the inclusion criteria, were included in the study.

The exclusion criteria were:

- a history of colonoscopy procedures;
- inflammatory bowel diseases;
- hereditary polyposis syndrome;
- established CRC;
- a history of significant intestinal surgeries (intestinal resection, bariatric surgery, etc.), except for appendectomy;
- any contraindications for polypectomy;
- incorrect and poor bowel preparation according to Boston Bowel Preparation Scale (BBPS) of 0–1 in any of the three bowel segments;
- patients with standard contraindications to colonoscopy (including acute diverticulitis/suspected perforation);
- incomplete colonoscopy procedure (technical difficulties);
- women who are pregnant or breastfeeding;
- any acute illness up to the time of inclusion;
- Oncologic diseases diagnosed within the past 3 years or with specific treatment completed less than 3 years ago.
- chronic kidney disease, autoimmune diseases, HIV, viral hepatitis B or C infections;
- chronic alcohol consumption;
- use of antibacterial, probiotic, and immunosuppressive drugs, as well as glucocorticosteroids and proton pump inhibitors in the last two months;
- diarrhea in the last two weeks.

### Sample collection

Patients were recruited during the consultation and a colonoscopy examination was scheduled, based on the indicated medical needs. During this consultation, after giving informed consent, patients were given a fecal collection kit. The procedure for collecting fecal samples on the day before bowel preparation was explained to the patients. Upon arrival for the colonoscopy examination, the collected fecal samples were immediately placed in a freezer at −20°C. Only those patients who had collected stool samples before the use of bowel preparation for the colonoscopy were included in the study.

### Colonoscopy procedure

The examination methodology complied with recognized standards and requirements and was described in detail in the author’s previously published work (14). Colonoscopy examinations were performed with the Olympus EVIS EXERA III (CF-HQ190L/l) video colonoscope. All examinations were performed by a single endoscopist who had 9 years of experience and performs over 1300 colonoscopies annually. In addition, colonoscopy procedures were performed under the supervision of an anesthesiologist, using short-term intravenous sedation based on propofol. The dosage of medication was determined by the anesthesiologist.

Bowel cleanliness was assessed by an endoscopist using the BBPS (Boston Bowel Preparation Scale). Four subjects were excluded from the study due to inadequate bowel preparation, as indicated by a score of 0–1 in any of the three bowel sections. The time of evacuation of the instrument from the cecum for each performed colonoscopy was not less than 7 minutes and was monitored by the endoscopy assistant.

Any detected polyps were described in the colonoscopy report according to the Paris (15) and NICE (16) classifications; the location and size of each polyp in the colon were also specified.

Morphological analysis was performed for all detected or removed polyps. When a polyp was identified, at least two biopsy samples were taken prior to polypectomy. Polyps were resected using either the cold loop or diathermocoagulation technique (hot loop), depending on size. In cases of non-resectable lesions, multiple biopsies were taken, and patients were referred for surgical treatment.

### Morphological diagnosis of lesions found during colonoscopy

All removed polyps and specimens from biopsied lesions were transmitted to the Academic Histology laboratory (Riga, Latvia) for morphological diagnostics. All samples were analyzed by expert pathologists and characterized according to the World Health Organization criteria, depending on morphological characteristics (17). All lesions were described as serrated polyps and lesions, low-grade dysplasia (LGD), high-grade dysplasia (HGD), superficial submucosal invasive carcinoma (SM-s; <1000 μm of submucosal invasion) and deep submucosal invasive carcinoma (SM-d; ≥1000 μm of submucosal invasion). No traditional serrated adenoma (TAS), sessile serrated lesion with dysplasia (SSL-D), or unclassified serrated adenoma were found morphologically.

### Determining microbiome composition by metagenome sequencing

Microbial DNA was extracted from faeces provided by study participants using the FastDNA Spin Kit for Soil (MP Biomedicals). The amount of DNA extracted was assessed using the Qubit dsDNA HS Assay Kit reagent kit. The DNA samples were stored at minus 20°C in the restricted-access facilities of the Latvian Genome Centre.

### Metagenome sequencing

Libraries for metagenomic shotgun sequencing were prepared using MGIEasy Universal DNA Library Prep Kit (MGI Tech Co., Ltd.). The input of DNA was 300 ng. Preparation steps briefly: DNA shearing into 420 bp fragments by S220 focused-ultrasonicator (Covaris) followed by size selection using magnetic beads; end repair and A-tailing; adapter ligation followed by magnetic beads cleanup of adapter-ligated DNA; PCR amplification and cleanup of the product; quality control; denaturation; single strand circularization; enzymatic digestion; cleanup of enzymatic digestion product; quality control. The quality and quantity of the resulting libraries was determined with an Agilent Bioanalyzer 2100 and a Qubit® 2.0 fluorometer. Metagenome sequencing was performed using the DNBSEQ-G400RS sequencing platform with the reagent set DNBSEQ-G400RS High-throughput Sequencing Set (FCL PE150) (MGITechCo., Ltd.) according to manufacturer’s instructions, obtaining 20 million reads per sample.

### Data analysis

Read quality evaluation was performed with FastQC (18). Adapter cutting and read trimming was performed with fastp (0.20.0) by using default trimming parameters. Paired reads with a length of 100 bp or longer were retained for further data processing. Reads originating from the host were removed with bowtie2 (v2.3.5.1) using GRCh38 as a reference. Taxonomic classification was performed with Kraken 2.1.2 and Bracken 2.7 against UHGG database (version 2) using Kraken’s confidence threshold value of 0.1. HUMAnN 3.8 (19) was used to perform functional profiling.

### Statistical Analysis

Statistical analyses were performed using R Studio 4.4.1 (20). Depth normalization for alpha diversity calculation was performed by constructing a multinomial distribution from metagenomic read count table and drawing n samples from each distribution where n=101889 is the minimum number of brackens classified reads for sample with the lowest coverage. For each sample alpha diversity indices (Shannon, Simpson, inverse Simpson and Pielou’s evenness) were calculated and this process was repeated 10000 times in total. The mean values of the diversity indices across iterations were then used as the representative alpha diversity metrics for each sample. Aitchison’s distance was used to evaluate the beta diversity. Transformed taxonomic data obtained by centered log ratio transformation with scikit-bio 0.5.5 were used for the construction of principal component analysis biplot with scikit-learn 0.22.

Differential abundance was tested with limma 3.60.2. Samples with less than 100000 assigned metagenomic reads were removed. Metagenomic features with at least 100 reads present in at least 10% of samples were retained. The P-value of < 0.05 was considered statistically significant. In addition, differential abundance and functional data were analyzed using the R packageMaAsLin2 (21), with adenoma status, sex, BMI, smoking status, and the presence of gastrointestinal diseases as fixed effects, and run ID as a random effect. A default Q-value (FDR) of < 0.25 was considered statistically significant.

## Results

### Demographic data

A total of 146 individuals were recruited for this study, including 85 polyp-free individuals, and 61 with colorectal adenomas. During the study, 10 patients were excluded (3 patients with ulcerative colitis, 3 with colorectal cancer, 4 with poor bowel preparation). All 136 patients were divided into two groups based on the presence (42%, n=56) or the absence (58%, n=80) of colorectal adenomas. Patient characteristics are summarized in Table 1.

**Table 1.**
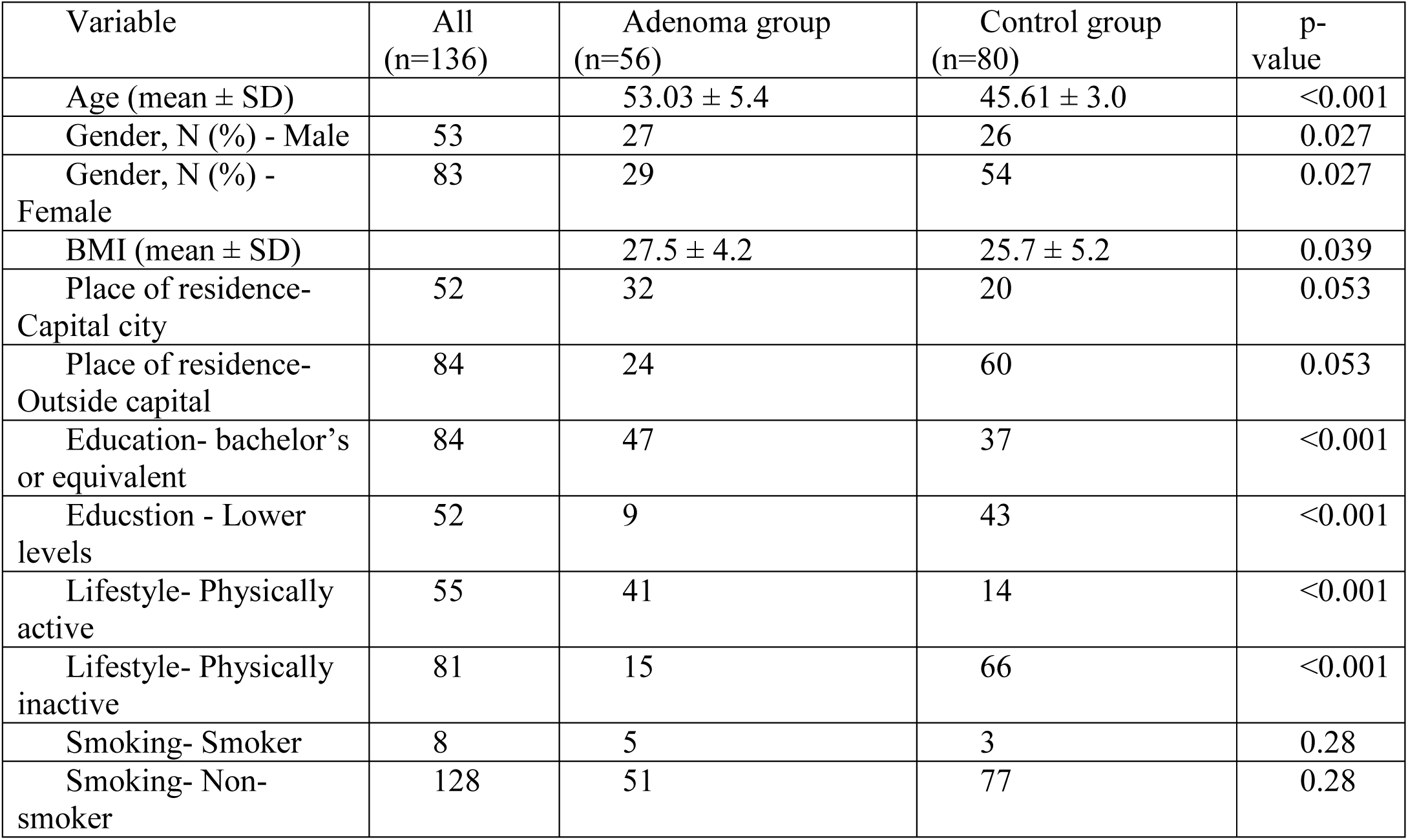
Demographic characteristics of adenoma and control group patients.

In total, 77 adenomas were found in 56 patients. From 56 patients 8 had high-risk adenomas (14%) and 48 had low-risk adenomas (86%), p<0.001.

Out of the adenoma-positive patients, 49% were male (27 patients) and 51% were female (29 patients), with a p-value of 0.05. The male/female ratio of the adenoma and control groups was 27/29 and 26/54, respectively. A total of 51.8% of female patients and 48.2% of male patients had at least one colorectal adenoma. The mean age of adenoma-positive patients was 55.3±13.5, while the mean age of adenoma-negative patients was 45.6±13 (p<0.001).

Among male patients, 15.4% of all detected colorectal adenomas were high-grade dysplasia (HGD), compared to 11.1% in female patients (p = 0.704). Overall, 41 patients (73.2%) had single tubular adenomas and 15 (26.8%) patients had multiple tubular adenomas (p<0.001).

The BMI of the adenoma group was 27.51 ± 4.18, while that of the control group was 25.73 ± 5.22 (p = 0.039). Statistically significant differences were found between patients with and without adenomas in age, gender distribution, educational level, place of residence, and physical activity habits (all p < 0.05). Patients in the adenoma group were older, more likely to live in the capital, more likely to have higher education, and significantly more likely to be physically active compared to the control group.

### Microbiome composition

The median number of paired-end reads obtained was 22830377 (IQR 10336229). After quality control and exclusion of host reads, 123 samples from 136 were retained for analysis (50 from adenoma positive and 73 from adenoma-negative patients), with a median of 22658664 reads per sample (IQR-interquartile range: 10326052). The median percentage of classified reads was 88.76% (IQR 2.71%).

The relative abundance of the detected genera in patients with colorectal adenomas (AD) and patients without adenomas (CO) is shown in Figure 1. The most common genera in both groups were *Faecalibacterium, Bacteroides, Blautia_A, Alistipes, Prevotella* and *Roseburia*. In addition, the archaeal representative *Methanomassiliicoccus_A* was found to be present among the 20 most common genera.

**Figure 1.**
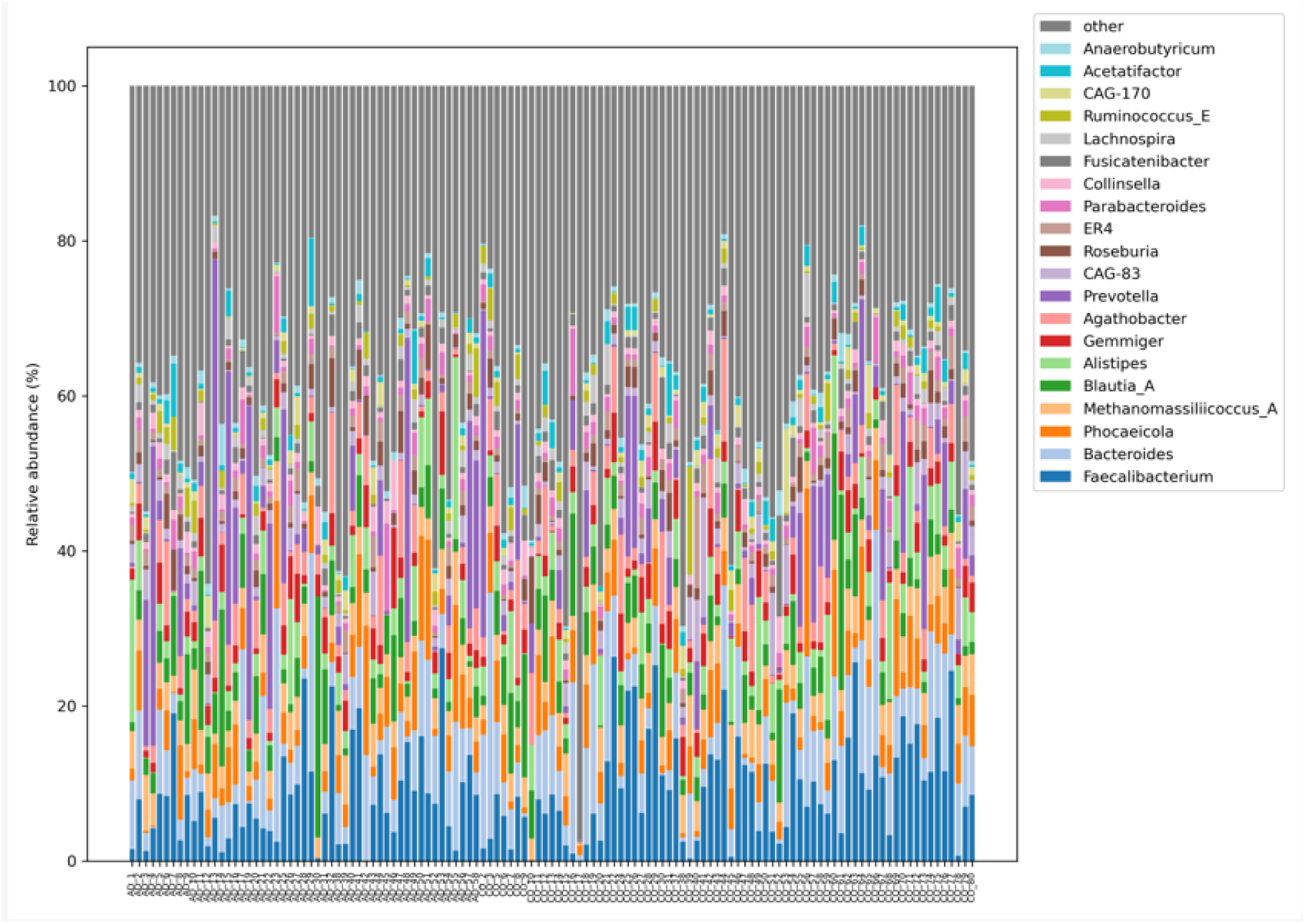
Genus-level taxonomic composition of fecal microbiota based on shotgun metagenomic sequencing

### Diversity analysis

#### Alpha diversity

Four commonly used diversity metrics were calculated: Shannon index, Simpson index, inverse Simpson index, and Pielou’s evenness. Data are presented as mean ± standard deviation (SD). Overall, alpha diversity showed slightly higher diversity in the control group (CO) compared to the adenoma group (AD), but none of these differences reached statistical significance.

#### Shannon index

The mean-4,6 ± 0,42 in the CO group and 4,53 ± 0,43 in the AD group.

#### Inverse Simpson index

The mean-41,47 ± 16,28 in CO and 37,68 ± 17,37 in AD.

#### Pielou’s evenness

The mean-0,65 ± 0,05 in CO and 0,64 ± 0,05 in AD.

#### Simpson index

The mean-0,97 ± 0.02 in CO and 0,97 ± 0,02 in AD. These results indicate a trend to higher microbial diversity in patients without colorectal adenomas; however, the observed differences were not statistically significant.

#### Beta diversity

Beta diversity analysis at the species level using PCA (Principal Component Analysis) showed statistically significant differences between AD and CO (p = 0.0002). The principal components PC1 and PC2 together explained 18.37% of the variance, indicating differences in the structure of the microbial community (Figure 2). The AD group had greater microbiota heterogeneity and a stronger association with the genera *Prevotella, Bacteroides, Collinsella* and *Holdemanella*, while the CO group was enriched with *Faecalibacterium*, *Anaerostipes* and other beneficial taxa. CO samples clustered more tightly, indicating a more stable microbiome.

**Figure 2.**
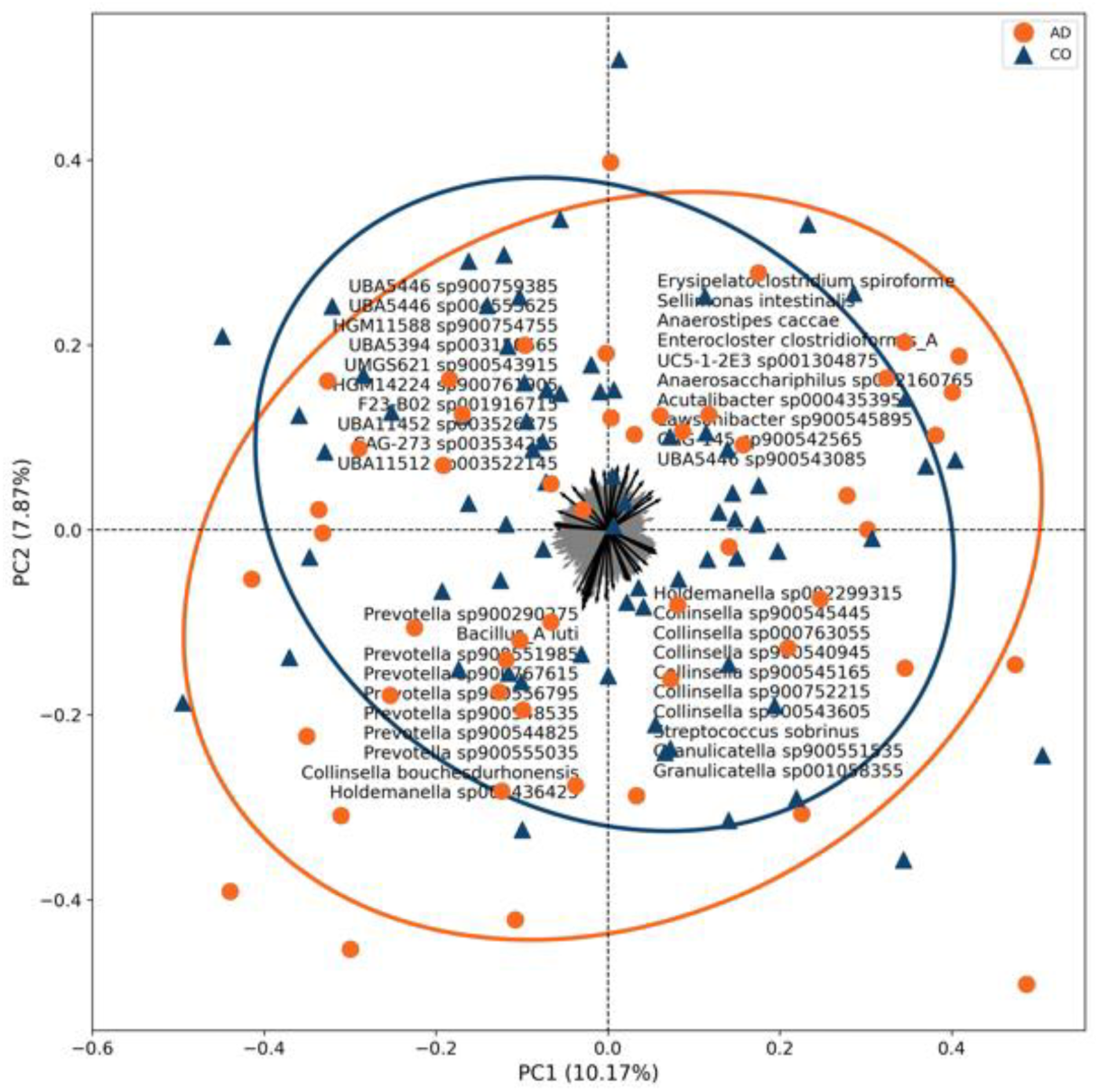
Principal component analysis (PCA) of beta diversity at species level in adenoma (AD, n= 50) and control (CO, n=73) groups

### Differential abundance analysis

Differential abundance analysis using MaAsLin2 with FDR correction (<0.25) identified one statistically significant altered taxon-*UBA7597 sp003448195*. This microorganism showed a significantly higher relative abundance in patients with adenomas compared to the control group (LogFC= 3.44; FDR= 0.002). A positive logFC value corresponds to a more than tenfold increase in the AD group. Although the mean expression value was low, the result reached strong statistical significance (p= 1.03×10⁻⁶), and the Bayes factor (B= 4.68) supported the reliability of the finding.

No other taxa were found to have significant differences between groups after FDR correction. Results were obtained by analyzing 123 metagenomic samples with the linear model approach (limma) and MaAsLin2 (Figure 3).

**Figure 3.**
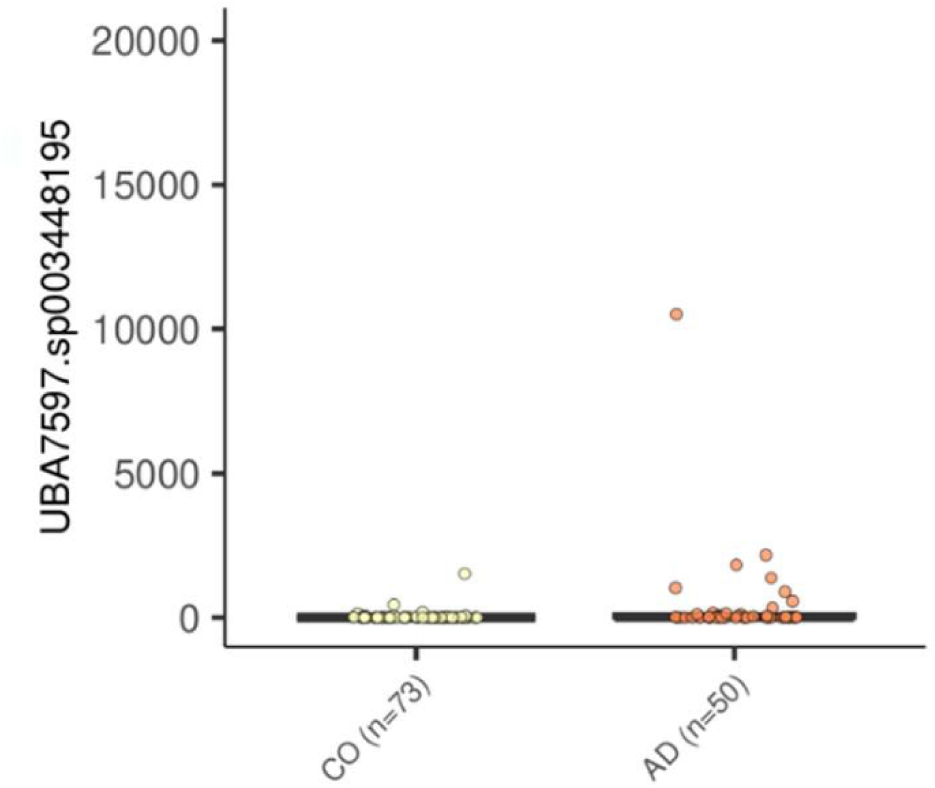
Differential abundance of *UBA7597 sp003448195* in adenoma (AD, n = 50) and control (CO, n = 73) groups identified by MaAsLin2 analysis

### Functional analysis

To investigate the potential differences in microbial functions depending on colorectal adenoma presence status, we evaluated the functional profile of the gut microbiome. Multivariable association analysis considering various fixed effects, revealed significant differences in six functions depending on colorectal adenoma presence using the MaAsLin2 model with an FDR <0.25. The results of the analysis have been summarized in Figure 4. Of the 136 participants included in the overall taxonomic analysis, one sample was excluded from the functional analysis (n = 135) due to insufficient quality or depth of sequencing data.

**Figure 4.**
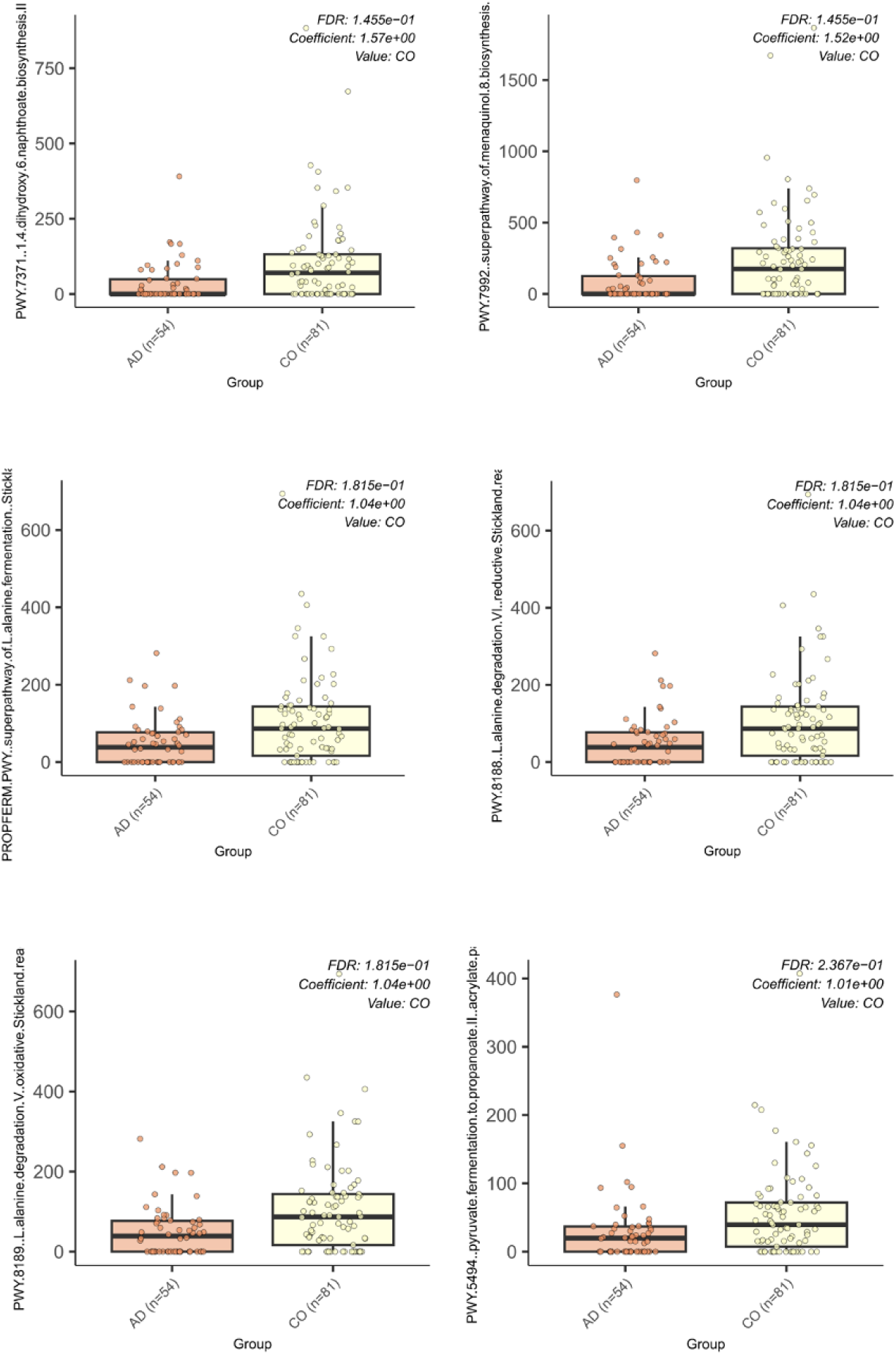
Differentially abundant microbial metabolic pathways between adenoma (AD, n = 54) and control (CO, n = 81) groups identified by MaAsLin2 analysis.

All pathways showed a higher relative abundance in the CO group. Two of them-PWY-7371 and PWY-7992-were associated with menaquinone (vitamin K2) biosynthesis (coefficients 1.57 and 1.52; FDR= 0.14). Three pathways - PROPFERM-PWY, PWY-8188 and PWY-8189 - corresponded to Stickland fermentation pathways with oxidative and reductive reactions of amino acids (each with a coefficient of 1.04; FDR= 0.18). In addition to PWY-5494 (pyruvate fermentation to propanoate II), the pathway that contributes to the production of SCFA (propionate) was also dysregulated in the AD group (coefficient 1.01; FDR= 0.23). All identified pathways were more widely distributed and had higher mean values in the CO group, while their reduced abundance and narrower distribution were observed in the AD group. Results were obtained by analyzing 135 metagenomic samples, and each of the pathways demonstrated differences between groups.

## Discussion

CRC is a major health concern due to its high incidence and mortality rates globally. The gut microbiota has emerged as a pivotal factor in CRC pathogenesis, with microbial dysbiosis being linked to colorectal adenomas, the precursors to CRC (22).

Despite considerable research (11–13), the relationship between gut microbiota and colorectal adenomas remains not fully understood. Previous studies (11–13) have provided inconclusive results, highlighting the need for further investigation into the microbiota composition of individuals with and without colorectal adenomas.

We evaluated the composition of the gut microbiota of 136 patients undergoing colonoscopy, dividing them into two groups based on the presence or absence of colorectal adenomas. Our results showed differences in the relative abundance of one microbial taxa between the two groups. In our study, most of the analyzed adenomas were LGD (tubular adenomas smaller than 10 mm without high-grade dysplasia and/ or villous structure. So far, a small number of publications have been published, where low-risk adenomas have been studied (23).

Colorectal adenoma incidence in our study appeared to be influenced by several host-related and environmental factors known to affect gut microbiota composition. Advanced age is associated with microbiota shifts toward a pro-inflammatory profile and reduced functional capacity, potentially contributing to neoplastic transformation (24–27). Hormonal and immunological differences between sexes may also modulate microbial communities and colorectal neoplasia risk (28, 29). In addition, urban living and higher educational level-typically linked to improved health awareness - can reflect lifestyle patterns such as lower fiber intake and higher consumption of processed foods, which are known to disrupt microbial balance and may promote adenoma development. The literature describes the effects of smoking on the composition of the microbiota, such as reduced diversity and a decrease in SCFA-producing bacteria (30–33).

### Differences in diversity

Previous studies on gut microbiota diversity in colorectal cancer have shown mixed results. Some studies report reduced diversity in cancer patients, while others find no significant differences. Our findings align with studies that suggest in alpha diversity alone may not be a sufficient indicator of colorectal adenoma presence (34, 35).

High diversity in the gut microbiome is generally associated with a healthy and resilient microbial community. The similar diversity indices in both patient groups suggest that factors other than microbial diversity such as specific microbial taxa or functional capabilities of the microbiome might be more critical in the development of colorectal adenomas (36, 37). Although alpha diversity scores were not significantly different, beta diversity analysis revealed that the overall microbial community structure was altered in patients with colorectal adenomas. This observation also reflects previous studies that have shown that individuals with colorectal adenomas have increased inter-individual microbiota variability and a shift away from the stable microbial profile (4). This pattern is consistent with findings by Deng et al. (36), who reported significant differences in beta diversity between patients with colorectal adenomas and healthy controls despite no significant changes in alpha diversity.

The genera *Prevotella*, *Collinsella*, and *Holdemanella* appeared more frequently in samples from the adenoma group, but these differences were not statistically confirmed by differential abundance analysis and were not clearly seen in the biplot, suggesting that their potential association with adenomas remains inconclusive. *Prevotella* is associated with chronic inflammation and mucosal immune activation, especially in Western populations, where a low-fiber diet has become prevalent in recent years (38–40).

The results suggest distinct microbial communities between the AD and CO groups, with specific microbial taxa characterizing each group. *Prevotella* is often associated with carbohydrate-rich diets and has been linked to inflammatory conditions in the gut (41). Specific taxa such as *Prevotella*, *Faecalibacterium* and *Bacteroides* can influence the development and protection against colorectal adenomas (42).

*Prevotella’s* association with inflammation and *Faecalibacterium’s* anti-inflammatory properties highlight the complex interactions between diet, microbiota, and gut health through butyrate production (38, 43, 44). Butyrate strengthens the gut barrier and modulates the immune response, potentially protecting against colorectal adenomas (45–48).

*Collinsella* is also known to affect intestinal permeability and bile acid metabolism and has been associated with metabolic disorders and low-grade inflammation (49, 50). The increase in the genus *Holdemanella* may reflect broader shifts in the composition of anaerobic microorganisms.

*Faecalibacterium* and *Anaerostipes*, which produce butyrate, a major short-chain fatty acid with described anti-inflammatory properties, were more abundant in the control group. *Faecalibacterium prausnitzii* has been extensively studied for its ability to maintain mucosal integrity, reduce oxidative stress, and induce regulatory T cells (43, 44, 51). Decreased abundance of this microorganism has been associated with the development of inflammatory bowel disease and CRC, making it a potential protective biomarker.

### Differentially abundant species between groups

Differential abundance analysis revealed one statistically significant taxon, *UBA7597 sp003448195*, which was more abundant in patients with adenomas. This taxon belongs to a relatively uncharacterized clade of intestinal bacteria and has not been extensively studied in the context of colorectal neoplasia. However, its increase in the adenoma group and statistical significance suggest a potential role in dysbiosis or mucosal damage. Although its biological function remains unknown, this finding highlights the need for deeper taxonomic and functional characterization of understudied microbial taxa in CRC studies (52). Deng et al. (36) reported higher abundance of *Esherichia, Shigella* and *Clostridium* species in polyp patients, supporting the notion that shifts in opportunistic or pathogenic microorganisms may play a role in early colorectal carcinogenesis.

### Comparative analysis of gut microbial functional profiles between adenoma and control cohorts

To complement the taxonomic analysis, we also assessed the functional differences in microbial metabolic pathways between groups. Six metabolic pathways showed significantly higher activity in the control group. Two of them were related to menaquinone (vitamin K₂) biosynthesis (PWY-7371 and PWY-7992), three were involved in Stickland fermentation (PWY-8188, PWY-8189 and PROPFERM-PWY) and one in SCFA production (PWY-5494 – pyruvate fermentation to propionate II).

The key bacteria of 1,4-dihydroxy-6-naphthoate biosynthesis II are *E. coli*, a well-characterized model organism for menaquinone biosynthesis (53), *Bacteroides fragilis*, which is involved in vitamin K2 production (54) and *Clostridium spp*., known for its role in anaerobic respiration and vitamin K2 biosynthesis. The key bacteria of Superpathway of menaquinol-8 biosynthesis are *Escherichia coli K-12*, *Klebsiella aerogenes*, which produces menaquinones for electron transport and *Bacillus subtilis*. Some *Lactobacillus* species can produce menaquinones, contributing to gut health (55).

Both of those functions are involved in pathway I of menaquinol-8 biosynthesis. 1,4-dihydroxy-6-naphthoate is a precursor for the biosynthesis of demethylmenaquinol-8 (56).

MK and DMK are crucial components of the electron transport chain in many bacteria, facilitating electron transfer between hydrogenases and cytochromes. They are especially crucial in anaerobic and facultatively anaerobic bacteria, contributing to processes like sporulation and cytochrome regulation. The conversion of this compound to DMK-8 occurs in several types of bacteria, particularly those involved in anaerobic respiration and vitamin K2 biosynthesis. *Clostridium* species are involved in diverse metabolic pathways, including the synthesis of menaquinones (57).

Reduced activity of menaquinone biosynthetic pathways in patients with colonic adenomas may be clinically relevant, as vitamin K2 is known to suppress proliferation and promote apoptosis in colorectal cancer cells (55, 58–60). The ability of bacteria to synthesize menaquinone is largely dependent on symbionts, such as *Lactococcus*, *Bacteroides*, and *Eubacterium* - taxa that are sensitive to dietary interventions and the effects of inflammation (57). Reduced vitamin K2 synthesis could contribute to cell cycle dysregulation and tumorigenesis in the colonic mucosa.

Previous findings (57, 61) support the idea that early changes in gut microbiota and their functions can precede and potentially contribute to the formation of colorectal adenomas, even when they are of low risk and small size. This is supported in Article by Deng et al. (36), who found microbial alterations in patients with tubular adenomas, which are known as pre-cancerous lesions, pointing the value of microbiota shifts as potential early biomarkers. It could be explained by changes in gut microbiota composition that occur early in adenoma development, even before they become high-risk or large. Early interactions between host cells and microbiota can initiate molecular changes contributing to adenoma formation.

Patients with adenomas also had reduced Stickland fermentation pathways, which are known to play a role in the oxidation and reduction of amino acids by anaerobic gut bacteria. These pathways play a role in nitrogen metabolism (62). Although their impact on carcinogenesis remains unclear, it is known that reduced Stickland fermentation activity may reflect a broader dysfunction of anaerobic metabolism in a dysbiotic microbiome.

The process of SCFA production pathway involves pyruvate being converted into propionate, which is then further metabolized into various products, including SCFAs. The key bacteria of Superpathway of L-alanine fermentation (Stickland reaction) are *Faecalibacterium prausnitzii, Eubacterium rectale* and *Anaerotignum propionicum*, which are known to participate in the fermentation of amino acids, including L-alanine, through pathways that produce short-chain fatty acids like propionate. *Anaerotignum propionicum* have the ability to ferment L-alanine to ammonia, CO2, acetate and propionate (63). It is known that *Anaerotignum faecicola* can ferment carbohydrates and produce SCFAs, including propionate production by gut bacteria, including species related to *Anaerotignum* (34). *Duncan et al*. highlighted the metabolic capabilities of related species in producing SCFAs (35). These data could explain the reason why colorectal adenomas are more common in the patient group where *Anaerotignum faecicola* is found less.

Finally, the activity of the PWY-5494 pathway was also reduced in the adenoma group. This pathway is involved in the synthesis of propionate, a short-chain fatty acid with known antiproliferative and immunomodulatory effects. Propionate and butyrate act synergistically to support epithelial barrier integrity and modulate gene expression in the colonic mucosa (64, 65). Lower activity of this pathway may impair the availability of short-chain fatty acids, promoting pro-oncogenic processes such as epithelial proliferation, immune dysregulation, and oxidative stress.

These findings suggest that the development of colorectal adenomas is associated not only with taxonomic shifts towards to genera with potential pro-inflammatory activity, but also with functional disruptions in key microbial metabolic pathways. Reduced vitamin K₂ biosynthesis, impaired SCFA synthesis capacity, and increased abundance of poorly characterized taxa such as *UBA7597* highlight the complex ecological and functional changes of the gut microbiome in the development of precancerous conditions.

Studying the types of microorganisms involved in metabolic processes and their interactions with gut bacteria could provide valuable insights into maintaining a healthy balance and preventing colorectal adenomas. Our research highlights the significance of these processes, in gut health and their potential role in preventing colorectal adenomas. By focusing on them we might come up with ways to support a gut environment and lower the risk of colorectal cancer. This study adds to existing data by providing metagenomic evidence, highlighting specific patterns with interesting and novel findings, and focusing on specific microbiome changes. Although this was a single-center study with a moderate sample size, its strengths include detailed colonoscopy procedures, strict exclusion criteria, and high sequencing depth. Interventions like probiotics or dietary changes that promote target bacteria could offer an approach to stopping the development of colorectal adenomas. The reduction in superpathway of menaquinol-8 biosynthesis among patients with adenomas emphasizes the importance of maintaining a balanced gut microbiome for preventing colorectal cancer.

Our study provides further evidence that the composition of the gut microbiota differs significantly between individuals with and without colorectal adenomas, and that this difference may be influenced by various demographic, lifestyle, and functional microbial characteristics. Our findings are consistent with a growing count of literature highlighting the role of gut dysbiosis in colorectal carcinogenesis, particularly in the early stages of the adenoma-carcinoma pathway.

Further studies using longitudinal designs, functional analyses, and strain-level resolution are needed to confirm the causality of these microbiome changes and understand their potential utility as biomarkers or therapeutic targets in colorectal cancer prevention.

Our study has several strengths that contribute to its reliability. Firstly, we used shotgun metagenome sequencing, which offers comprehensive insights into the microbial communities of the gut. This method provides a detailed analysis of both-the taxonomic composition and functional potential of the microbiome. Secondly, our study included well-characterized patients, ensuring accurate and relevant data collection. The single-center case-control design, conducted by an experienced endoscopist with an appropriate and sufficient adenoma detection rate enhances the validity of our findings. Additionally, detailed exclusion criteria minimized potential confounding factors, thereby increasing the precision of our results.

Limitations of our study include a young patient group and a small sample size. Additionally, we did not examine microbiome structures across different populations.

## Conclusions

The identification of microbial taxa and functional pathways linked to adenoma presence highlights the potential for using microbiota-based markers and treatments to prevent and manage colorectal cancer. Future research should focus on establishing connections and clarifying how gut microbiota influences colorectal carcinogenesis.

## Author contributions

all authors made substantial contributions to the manuscript following the guidelines set by the International Committee of Medical Journal Editors (ICMJE). Individual contributions are detailed below:

**Ilona Vilkoite**: conceptualization and design of the study, data acquisition, interpretation of the results, manuscript drafting, and revisions, collection and analysis of fecal microbiome data, recruitment of patients, clinical data collection, revising the manuscript.

**Laila Silamiķele:** expertise in microbiome analysis, data interpretation, and review for intellectual content, manuscript drafting, and revision.

**Jānis Kloviņš, Ivars Tolmanis, Aivars Lejnieks:** overall project, study design and methodology, review of the final version.

**Ivars Silamiķelis, Elīna Runce, Ksenija Margole, Olga Sjomina, Krista Cēbere**: statistical analysis, data visualization, substantial contributions to the manuscript’s revision process.

**Conflicts of Interest:** the authors declare no conflict of interest.

**Data availability statement:** the data acquired in the study are deposited in the European Nucleotide Archive (ENA), accession number PRJEB79034

## Acknowledgments

We thank the Genome Database of the Latvian Population for their support in processing the biological samples and related data used in this study.

